# Local graph-motif features improve gene interaction network prediction

**DOI:** 10.1101/2025.02.21.639582

**Authors:** Victor Julio Leon, Jordan K. Matelsky, Amanda Ernlund, Lindsey M. Kitchell, Kristopher D. Rawls, Caitlyn Bishop, Elizabeth Reilly

## Abstract

Gene interaction networks specify how genes interact to produce an organism’s phenotype. These networks are often incomplete due to absent or unobserved information. Predicting these missing links is critical for many applications, including genome-wide association studies and phenotype prediction. Efforts have previously applied graph neural networks (GNNs) to this missing-link prediction problem, but these techniques too have limitations when the sparsity of the networks is very high. Here, we apply a novel feature engineering technique that uses local graph motif incidence to enhance the feature set for variational graph autoencoders (VGAE). We compare the performance of our technique against state-of-the-art approaches, and then progressively hide more and more of the original graph edges. Our results show that VGAEs with our local-area motif prevalence (LAMP) features outperform state-of-the-art node embeddings for a wide range of missing edges on both a benchmark and a biological dataset. We also observe that this combined VGAE and LAMP technique has the potential to facilitate the search for novel genetic interactions in an experimental adaptive sampling context with far fewer samples. Improvements to gene interaction imputation can lower the barrier to new pharmaceutical and epidemiological discoveries by revealing hidden gene interactions that steer the development of potential drug targets.

## Introduction

Gene interaction networks are graphical representations of how an organism’s genome interacts to produce its traits, or phenotypes^1^. The networks are generally represented as a graph *G* = ⟨*V, E, A*⟩ where *V* is a set of vertices each representing a gene, (*u, v*) ∈ *E* is a set of edges where *u, v* ∈ *V* represent an interaction between *u* and *v*, and *A* is an optional set of attributes (such as gene names or molecular metadata) that can be assigned to either vertices or edges. In rare cases, the gene interaction network of a species is well-studied and well-known. More commonly, however, gene interactions are sparsely characterized, and thus the network suffers from many unknown edges^2^.

Genome-wide association studies (GWAS) and phenotype prediction are two of the many applications in the biology domain which require complete gene interaction networks. These techniques are now commonly applied to crop evaluation, epidemiology, and pharmacology^3,4^. Many hopeful studies, however, are stymied by datasets where many or most of the edges denoting gene interactions are unknown. Thus, edge prediction in gene interaction networks is a timely and important capability.

Recent work has demonstrated the promise of machine learning methods in addressing the challenge of edge prediction in gene interaction networks. The most common methods use a variety of neural network architectures, such as deep variational autoencoders^5,6^. Despite ongoing progress in gene interaction prediction, many techniques still fail to consider the rich, underlying graph structure of the data^7^. More recently, methods based on graph neural network (GNN) architectures have been proposed for gene expression imputation. GNNs perform feature extraction by exploring the local neighborhood of a node in the gene interaction network in order to produce features for each vertex^7,8^. Specifically, variational *graph* autoencoders (VGAE) have been shown to help address this imputation task, likely due to the improved graph context of these models^2,8,9^. Using higher-order graph features and vertex or edge attributes is a promising approach to improve the performance of these models.

In this work we propose a motif-based feature extraction technique that uses the set of subgraph instances to which a vertex belongs to produce a unique embedding. In our technique, we enumerate instances of selected subgraph motifs within the interaction graph and use these explicit subgraph counts to enhance the gene expression network node feature set (**Fig. 1**). Using the DotMotif network-motif search algorithm, we are able to efficiently and precisely count the number of times a given motif appears in the local neighborhood of each vertex in the graph, in a highly parallelizable manner^10^. We evaluate our technique’s performance on a VGAE^11^ and discover that we perform at or above state-of-the art in a link prediction task. We progressively remove edges from the partial starting-graph and observe that our technique outperforms state-of-the-art for edge-imputation across a wide range of missing edges on both a benchmark (FB15k-237)^12,13^ and a biological dataset of a eukaroytic cell^1^. This scalable approach is applicable to graphs of various sizes, as the motif finding process is highly parallelizable and can be sparsely subsampled to improve performance, at the cost of feature accuracy. This work has the potential to facilitate the search for novel genetic interactions in an experimental adaptive sampling context with far fewer samples than competing techniques. Furthermore, our approach is applicable to a wide range of graph-based machine learning problems outside the field of gene interaction, including social network analysis and chemical compound prediction.

**Figure 1.**
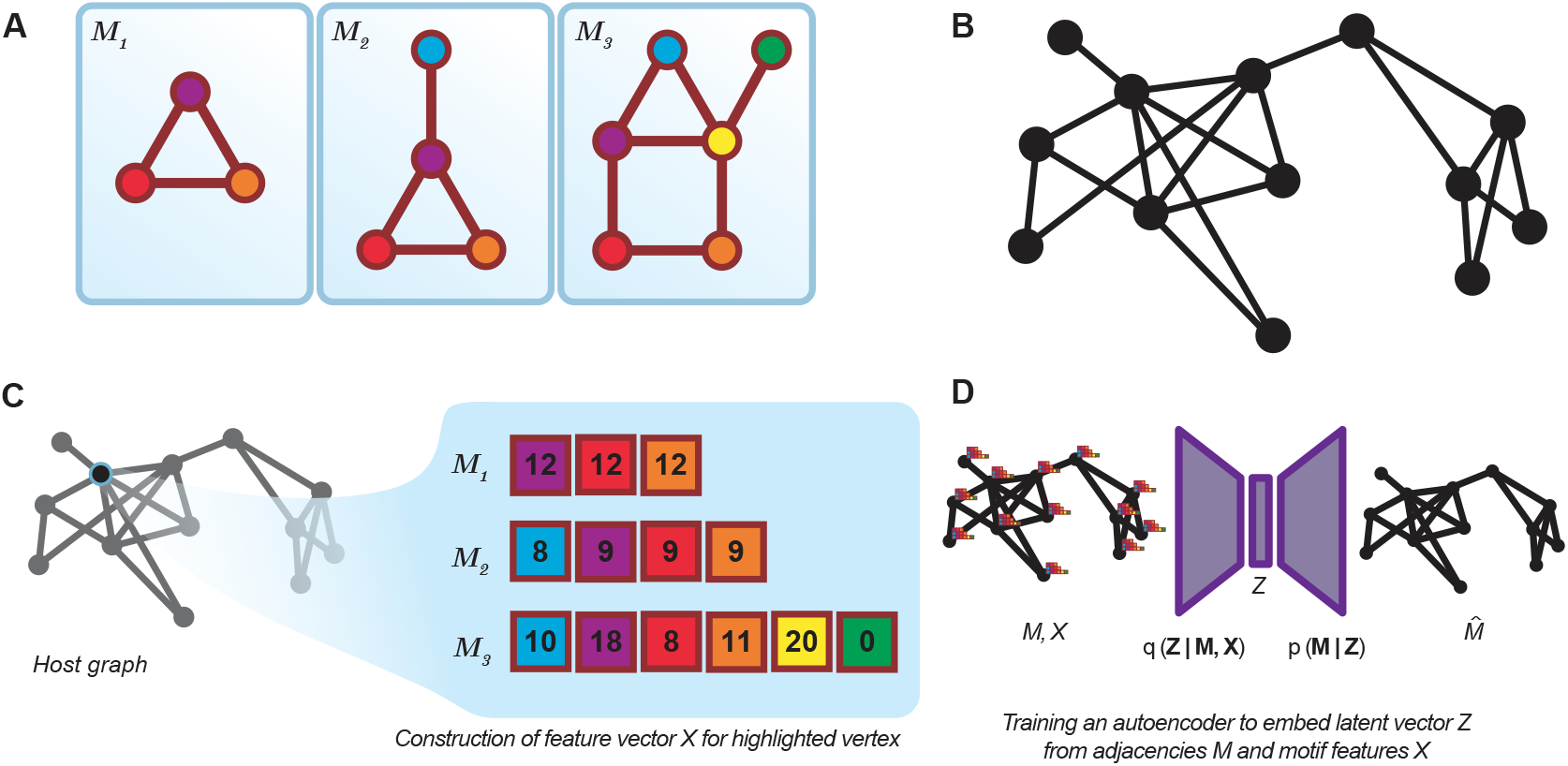
A visual overview of the LAMP algorithm. A. An example of three motifs used for LAMP feature generation. These motifs were hand-selected to have minimal automorphisms. **B**. An example host network. This network is undirected, though LAMP works on directed graphs as well as multigraphs. **C**. For a given vertex in the host graph (black, left), local motif participation is enumerated. Because this vertex participates as the “antenna” of the motif *M*_2_ in eight different ways, the first element of the feature vector (cyan square) is 8. These per-motif feature vectors are concatenated and fed into the downstream graph neural network. **D**. A schematic of the VGAE architecture, illustrating a graph enriched with LAMP features *X* prior to encoding in the encoder *q*.

## Background

Our proposed approach combines knowledge of gene interaction networks, variational graph autoencoders, and graph motif search. In this section we will introduce these topics and comment on strategies for marrying these technologies to improve gene interaction imputation and edge imputation more broadly.

### Gene interaction networks

Gene interaction networks are formalized as a graph *G* = ⟨*V, E, A*⟩, where nodes / vertices *V* represent genes, and edges (*u* ∈ *V, v* ∈ *V*) ∈*E* represent (undirected) interactions between genes. ^1^ These edges can represent interactions of various types, including physical interactions, genetic interactions, and regulatory interactions. Different types of interactions can be represented by adding edge or vertex attributes *A* to the graph and the library of motifs, though we do not address these in this work.

Gene interaction graphs are generated through a variety of experimental techniques, including yeast two-hybrid assays, synthetic genetic array analysis, and other analyses^14,15^. Due to the cost and complexity of these experiments, gene interaction networks are often sparsely characterized, and thus contain many unknown-unknown edges. Because these graphs are binary and undirected, it is difficult to draw a clear line between the case of an unexplored or unobserved edge and the case of a non-interacting pair of genes. For this reason, gene interaction graph edge imputation is a challenging problem in lesser-studied organisms and in the case of large-scale experiments.

### Variational Graph Autoencoders

Graph variational autoencoders (VGAEs) are a class of graph neural network (GNN) that use a variational autoencoder (VAE) architecture to learn a low-dimensional representation of a graph^11^ and have been proposed for a variety of graph-based machine learning tasks, including link prediction, node classification, and graph classification^7^. The basic architecture consists of an encoder that maps the graph into a low-dimensional latent space, and a decoder that reconstructs the graph from the latent space. The intention is that this low-dimensional representation will capture the essential structure of the graph, and that the decoder will be able to reconstruct the graph from this low-dimensional representation even when some information has been lost.

### Motifs and subgraph search

Like many other networks, such as social graphs or knowledge graphs, gene interaction networks are known to contain recurring, self-similar subgraphs^10,16^. The subgraphs that occur with statistically significant frequency — known as motifs — are thought to be responsible for many of the network’s emergent or repeating properties^16^, and it is thus of interest to identify those motifs in gene interaction networks where they may correspond to biologically interpretable or therapeutically useful mechanisms. These graph motifs are not to be confused with genetic motifs, which are short repeating DNA or RNA sequences that are known to have a biological function^17^.

Motif enumeration can also be used to reduce the dimensionality of a graph, by counting the number of times a given motif appears in the local neighborhood of each vertex in the graph. In GNNs, such an embedding can be performed as a preprocessing transformation^18–20^, or as a derived, learnable property itself^21^.

DotMotif is a graph motif search algorithm that uses an induced monomorphism search algorithm to find motifs in a larger graph^10^. The algorithm is highly parallelizable and can be used to find motifs in large graphs, such as gene interaction networks, in a reasonable amount of time — especially if constraints are placed on the size or attributes of the motifs to be found.

Using the DotMotif algorithm, we can efficiently and precisely count the number of times a given motif appears in the local neighborhood of each vertex in the graph. We will use this information to enhance the feature set for our VGAE and compare the performance of our technique against *node2vec* and the Jaccard index, progressively hiding more and more of the original graph edges.

## Methods

Our technique comprises a novel feature engineering technique and we show a practical use of this feature engineering approach for missing-link prediction in a gene interaction network. We first describe the underlying algorithm for producing local-area motif participation features, and then describe the training and validation steps we used to compare this approach against the current industry best-practices.

### Local-area Motif Participation

We want to generate a set of features that describe the local graph neighborhood of a vertex in a concise but maximally unique vector “fingerprint.” It is well-known that subgraph monomorphisms well-describe the local properties of a graph^10^ and so we propose that a valuable feature vector to uniquely identify a vertex can be described by how that vertex participates in a small library of subgraphs. We call this feature set the local-area motif participation (LAMP) of a vertex (**Fig. 1**) which can be used to uniquely embed vertices of a target graph.

The first step of LAMP feature generation is the selection of motif library *M*, an unordered set of subgraphs (**Fig. 1A** that will be counted in the host network. In general, this set should be curated to be a small set of graphs for which no motif is a subgraph of any other motif (or at least such cases are minimized, as they result in a higher runtime cost without adding new information), and for which automorphism symmetries are as few as possible (as automorphisms will result in axes of the LAMP feature vector that are perfectly linearly correlated and thus yield no additional information). In general the selected motifs should be simple (i.e., fewer edges per motif will result in faster runtime): Runtime of subgraph monomorphism search scales exponentially with the number of edges in a motif. For this work, we arbitrarily selected a cycle, a cycle with a connected edge to reduce automorphic symmetry, and a combination of adjoining cycles (*M*_1_, *M*_2_, and *M*_3_ respectively in **Fig. 1A**), although similar performance characteristics were found anecdotally using other combinations.

Our proposed algorithm is described in Algorithm 2. The algorithm starts with a host graph *G* and a motif library *M*. For each vertex *v* ∈ *V*_*G*_ and each vertex 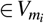 in a motif *m*_*i*_ ∈ *M*, we count the occurrences of the motif *m* and the number of times vertex *v* is mapped by the monomorphism function to vertex *u*. The process continues for each node in the graph. Note that this node-level motif search can be performed in parallel for any size graph, meaning that while the subgraph monomorphism algorithm is NP-hard, it can be run on a sufficiently small subset of the host graph that captures only the local graph area surrounding a vertex, which can in general reduce performance bottlenecks.

In this work, LAMP features were computed by searching for subgraph monomorphism instances of the library *M* in the host (dataset) graphs using the GrandIso/DotMotif software suite.^10^ The proposed induced monomorphism search can be carried out either at the vertex-level (i.e., searching only in the local neighborhood of the vertex as described above), and thus was parallelized across vertices. Alternatively, an exhaustive motif search can be run globally on the entire host network and then post-processed. The latter technique is slightly faster, if hardware can support storing the complete search in memory at once. The size- and time-complexity characteristics of the GrandIso algorithm are described in Matelsky et al., 2021^10^.

### Datasets

LAMP, node2vec, and the Jaccard index are compared on the benchmark FB15k-237 dataset, which has 14,541 nodes and 272,115 edges^12,13^. Then, we measure the models’ performance on a biologically relevant, comprehensive genetic interaction network of a eukaryotic cell with 5,362 genes (nodes) and 25,405 interactions (edges)^1^. Herein, we study undirected graphs. We follow the data cleaning procedure described in the Supplementary Materials of Costanzo et al.^1^ with a Pearson correlation coefficient >0.2 indicating a genetic interaction.

#### Algorithm 2 Pseudocode for the LAMP feature extraction algorithm. The for-loops can be parallelized across vertices.

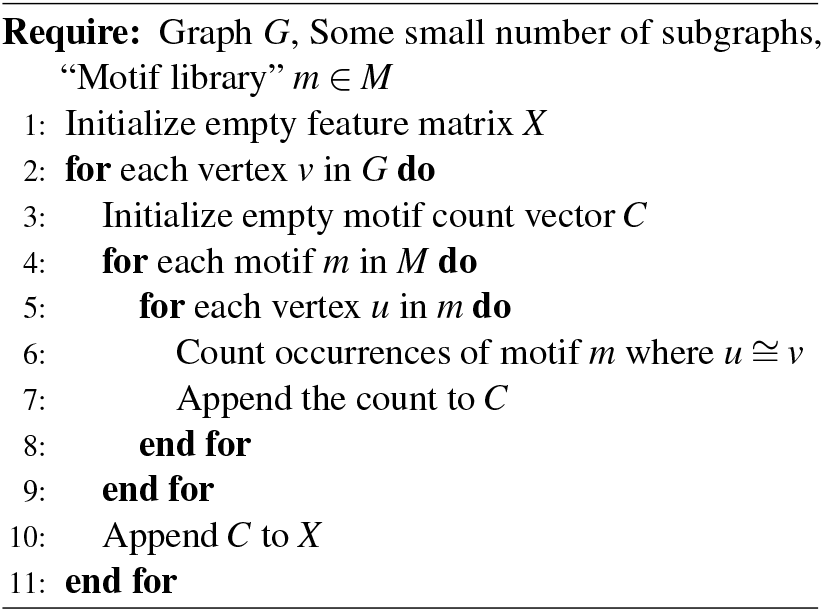

### Training and evaluation

In this work, we compare the performance of a VGAE trained with LAMP features (LAMP) against a VGAE trained with node2vec features (node2vec)^22^ and the Jaccard Index^22,23^. VGAE and node2vec are used since they perform well compared to other state-of-the-art models^2,24–26^. The Jaccard index is included as it is a common heuristic used in link prediction tasks^2,23^. The amount of edges hidden to the feature generating algorithms is varied by randomly removing various amounts of the networks’ edges, with the goal of studying the performance of the various models over a wide range of graph densities (**Fig. 2**). After generating features, the set of edges not used during feature generation is split into a 80% train and 20% test split. Performance is evaluated with 3 random train-test splits.

**Figure 2.**
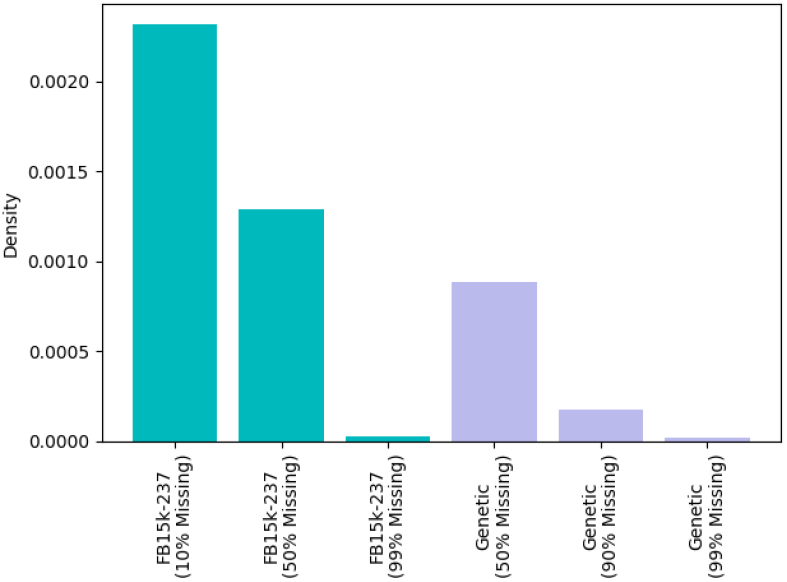
Network densities of subsampled graphs. When evaluating *node2vec* and *LAMP* features, only a subset of the edges are revealed to the feature-generation algorithm. In this figure we compare the network densities, computed as 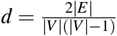.

Performance is measured using area under the receiver operating characteristic curve (AUC) and average precision (AP). Both datasets are quite sparse, with few edges between nodes. In the gene-interaction network, this can both represent the absence of a measurement or the confirmed absence of an interaction between genes. Since we are interested in predicting genetic interactions (positive samples), AP is the most relevant metric in this data-imbalanced case^2,24,25^. AUC also is included for completeness and ease of comparison with previous literature on link prediction. We evaluate model AP and AUC on balanced datasets, which has been noted in recent literature to over-estimate model performance^27^. In a balanced dataset, the train and test set have 1 non-edge for each edge.

More concretely, 10% missing edges during feature generation means 90% of edges are used to generate features (i.e. Jaccard indices, node2vec features, and LAMP features) and the remaining 10% of edges are used to train and evaluate the models (e.g. VGAE model). This means that, for the FB15k-237 dataset with approximately 272k total edges, the 50% missing edges condition during feature generation has 272k edges available for train and test of the VGAE (136k edges and 136k non-edges), while the 99% missing edges condition has 538k edges available for train and test of the VGAE (269k edges and 269k non-edge). The amount of edges available for training of the VGAE model sets the lower bound on % missing edges for each dataset. For example, we do not evaluate 10% missing edges condition for the genetic dataset because 1k edges is not enough to consistently train and evaluate a VGAE. We also study the effect of train and test set size on VGAE model performance by performing a study with constant 90% edges missing for feature generation while varying the train-test set size.

All VGAE models are trained with the ADAM optimizer^28^. For the FB15k-237 dataset with node2vec, we observe the best performance with 200 epochs and learning rate *η* = 1 × 10^−1^ for all missing edge percentages.

For LAMP with the FB15k-237 dataset, we observe the best performance with 200 epochs and learning rate *η* = 1 × 10^−2^ for both 10 and 50% missing edges and 200 epochs and learning rate *η* = 1 × 10^−1^ for 99% missing edges. For the genetic dataset, for both node2vec and LAMP and all missing edge amounts, we observe the best performance with 2000 epochs and learning rate *η* = 1 × 10^−2^.

An important parameter that we do not vary in this study are the motifs used for feature generation. We use the motifs in Figure 1 to provide a baseline of LAMP’s performance across two different graphs. We expect that LAMP feature performance would be influenced by choice of motifs.

We use the standard implementations of VGAE and node2vec in PyTorch Geometric with default hyperparameters^29^. NetworkX is used to calculate Jaccard coefficients^30^.

## Results

### Model evaluation

VGAE with LAMP features (LAMP) outperforms both Jaccard Index and VGAE with node2vec features (node2vec) for a wide range of missing edges for feature generation and training set sizes on the benchmark and biological datasets (**Tables 1 and 2**). The Jaccard index performs much worse than the other two methods across all proportions of edges missing and datasets.

**Table 1.** Performance of the three link prediction models on the benchmark FB15k-237 dataset. For a wide range of % edges missing for feature generation and training set sizes, LAMP has the highest AP and AUC. Clearly, as training set size increases, LAMP performance increases more significantly than the negative effect of higher % missing edges. At 10% missing edges, node2vec outperforms LAMP. We hypothesize that this is due to the smaller training set size and the relatively small motifs chose (3 to 7 edges), which may be too small to capture informative larger motifs in the graph with less missing edges. Each % edges missing is a unique subsample of edges from the overall graph. VGAE performance metrics are measured for 3 different train-test splits of graph edges with results presented as *mean* ± *standard deviation*. Jaccard index is deterministic for a given graph.

| % Edges Missing | Training Set Size | Method | AUC | AP |
| --- | --- | --- | --- | --- |
| 10 | 44k | Jaccard Index | 0.60 | 0.01416 |
|  |  | **node2vec** | **0.7588±0.0100** | **0.8172±0.0106** |
|  |  | LAMP | 0.5214±0.0125 | 0.5729±0.0128 |
| 50 | 218k | Jaccard Index | 0.54 | 0.001638 |
|  |  | node2vec | 0.7462±0.0327 | 0.8026±0.0199 |
|  |  | **LAMP** | **0.7932±0.0065** | **0.8334±0.0044** |
| 99 | 432k | Jaccard Index | 0.50 | $2.603 \times 10^{-3}$ |
|  |  | node2vec | 0.8390±0.0050 | 0.8682±0.0014 |
|  |  | **LAMP** | **0.8621±0.0014** | **0.8729±0.0016** |

**Table 2.** Performance of the three link prediction models on the comprehensive genetic interaction network of a eukaryotic cell. We observe that LAMP outperforms the other two models for a wide range of % missing edges for feature generation and training set sizes (i.e. 50 and 90 percent missing). This demonstrates that choosing motifs can be more informative than relying on node2vec’s random walk based featurization on a biologically relevant graph. On the higher ends (99% edges missing), node2vec features outperform LAMP features. The LAMP method is likely sensitive to the motifs chosen (Figure 1). At 99% edges missing, only around 250 edges are available for LAMP feature generation. In this case, there may not be enough edges to form the motifs selected for LAMP. node2vec is set with default 128 steps random walk, which can still take advantage of the lower number of missing edges. Each % edges missing is a unique subsample of edges from the overall graph. VGAE performance metrics are measured for 3 different train-test splits of graph edges with results presented as *mean*±*standard deviation*. Jaccard index is deterministic for a given graph.

| % Edges Missing | Training Set Size | Method | AUC | AP |
| --- | --- | --- | --- | --- |
| 50 | 20k | Jaccard Index | 0.66 | 0.03884 |
|  |  | node2vec | 0.8426±0.0070 | 0.8589±0.0051 |
|  |  | **LAMP** | **0.8519±0.0082** | **0.8762±0.0043** |
| 90 | 36k | Jaccard Index | 0.51 | 0.001849 |
|  |  | node2vec | 0.8292±0.0037 | 0.8502±0.0021 |
|  |  | **LAMP** | **0.8683±0.0032** | **0.8960±0.0037** |
| 99 | 40k | Jaccard Index | 0.50 | $2.603 \times 10^{-5}$ |
|  |  | **node2vec** | **0.7782±0.0015** | **0.7870±0.0005** |
|  |  | LAMP | 0.6896±0.0063 | 0.7154±0.0077 |

Counterintuitively, LAMP and node2vec performance is observed to increase with % edges missing for feature generation (e.g. from 10 to 99% missing edges for FB15k-237 and from 50 to 90% edges missing for the biological dataset). This is an artifact of the interaction between the amount of data available for feature generation and training set size on model performance. To isolate the effect of training set size on model performance, we vary train set size in the biological dataset for a constant 90% edges missing in Figure 3. As train set size increases LAMP model performance significantly increases, while node2vec performance stays relatively constant. LAMP features perform better than node2vec at larger train set sizes, implying that LAMP features are more informative than node2vec features. We hypothesize that LAMP’s worse performance at smaller train set sizes is due to having relatively few and sparser features vs node2vec (13 for LAMP vs 128 for node2vec), making the LAMP features we generated using 3 motifs more difficult to learn. Further studies on the effect of motif choice during LAMP feature generation could help improve LAMP feature performance at smaller train set sizes.

**Figure 3.**
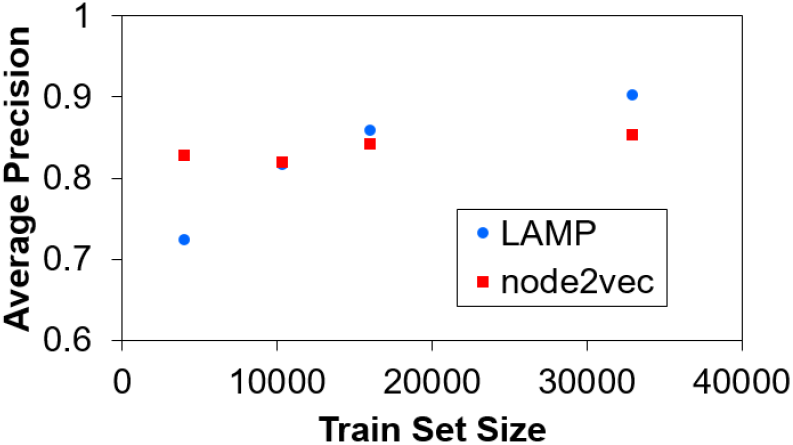
As train set size increases, model performance increases for LAMP and stays relatively constant for node2vec. Figure generated for VGAE with LAMP features generated with 90% of features missing for various training set sizes.

LAMP performance also drops below node2vec for 99% missing case on the biological graph. Here, both models show a reduction in performance relative to less % edges missing, but LAMP has a larger performance drop than node2vec. We hypothesize that this bigger drop in performance for LAMP is due to the motifs selected having 3 to 7 edges (Figure 1). At 99% missing edges in the biological graph, we only expect around 250 edges to be available for LAMP feature generation. At this point, LAMP features for motifs of 3 to 7 edges are very sparse. In contrast, node2vec is based on random walks, which is likely less sensitive to this case of very few edges available for feature generation.

Another area where node2vec outperforms LAMP is in terms of computational cost as number of edges increases. Motif feature generation takes significantly longer. This is because DotMotif runtime to identify motifs increases approximately exponentially with number of edges. One attractive strategy to make such a problem computationally tractable is to subsample the graph and generate features on the subgraphs. This strategy is particularly attractive because we observe that the performance of LAMP is robust for a wide range of % missing edges.

### Case study

The AUCs and APs reported in Table 2 and discussed thus far are useful to compare between predictive models since they are average model performance statistics. Here, we demonstrate how LAMP can significantly increase interaction sampling efficiency in practice using the 90% missing case of the biological genetic interaction network.

The activations output by LAMP for each edge in the balanced test set of 50% positive (4k samples, edge present) and 50% negative (4k samples, edge not present) are shown in Figure 5. In practice, a threshold activation is set by the user of the model to determine what the model predicts to be an edge or non-edge. For example, setting an activation threshold of 0.8 means that any edge with activation less than 0.8 is considered as a non-edge and any edge with activation greater than 0.8 is considered as an edge to sample. A higher or lower threshold is a trade-off between capturing all true edges (lower threshold) and minimizing how many non-edges are incorrectly predicted by the model to be edges (higher threshold). To evaluate the model in a more practical setting, here we report counts of true positives (TP), false positives (FP), false negatives (FN), precision, and recall for various thresholds.

LAMP has learned how to distinguish true edges from true non-edges, since true edges generally have higher activations and true non-edges generally have lower activations (Figure 5). A threshold of 0.55 leads to TP=3324, FP=1265, FN=789, precision=0.72, and recall=0.81. A threshold of 0.8 leads to TP=2409, FP=112, FN=1704, precision=0.96, and recall=0.59. Figure 4 summarizes the precision and recalls for various thresholds. The specific choice of a lower or higher threshold depends on the user’s preferred balance of precision and recall.

**Figure 4.**
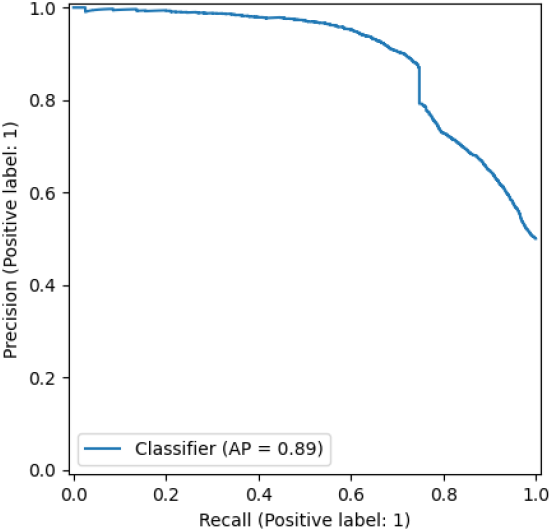
Precision-recall curve for LAMP with 90% missing edges in the biological dataset during feature generation.

**Figure 5.**
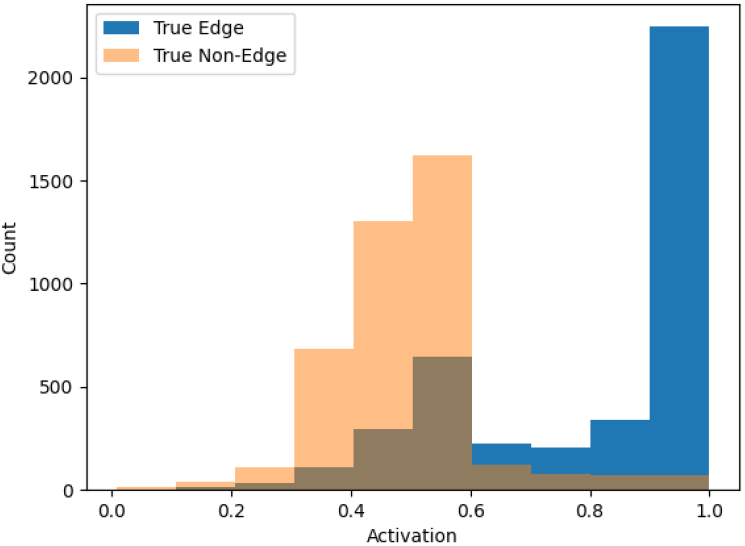
LAMP output activation on the test set. LAMP was trained on the 90% missing biological genetic interaction network.

Clearly, using LAMP with a 0.55 threshold is significantly better than random uniform sampling of interactions, which would yield, in this case of 50/50 interaction to non-interaction, a precision of 0.5. Actually, the performance of uniform random sampling on the genetic interaction graph would be significantly worse than 0.5, since the interaction graphs are low density (**Fig 2**).

## Discussion

In this paper, we propose an edge-imputation technique for gene interaction networks that marries the advantages of variational autoencoders (VAEs) with the structural advantages of graph neural networks through an architecture known as a Variational Graph Autoencoder (VGAE) network^7,11^. Furthermore, we enrich the network features with our novel algorithm that produces local-area motif participation (LAMP) features, which are generated using the DotMotif algorithm^10^. Our model serves as a strong predictor of local graph structure and can recover missing edges even in the very high-missingness regime.

We believe this exceptional performance is due to a more holistic encoding of the complex, high-dimensional structure of the input dataset. Though a deeper analysis is required to establish causal relationships between edge recovery rate and the semantic, biological significance of the recovered data, we believe that this work serves as a strong foundation for future work in the field of network edge imputation.

This approach offers researchers the promise to sample thousands of genetic interactions instead of tens of millions, as done in previous genetic interaction studies^1^. Furthermore, LAMP has the potential to facilitate the search for novel genetic interactions in an experimental adaptive sampling context with far fewer samples.

We anticipate that future work will include more deliberate motif library design; in this work, we were able to achieve high performance with hand curated graph motif sets but we expect that performance can be tuned even further. Indeed, the ideal motif set may be a function of the task or the dataset. A deeper study into the causes of the cross-over in Figure 3 could provide insight into motif selection. We also anticipate that future work will include the use of edge attributes in the motif search, as well as the use of more complex graph neural network architectures. The DotMotif motif search tool is already capable of handling complex vertex and edge attributes, and we anticipate that this will be a fruitful area of future work as attributed subgraph search can be dramatically faster and more informative than that of unattributed subgraphs. We also intend to apply this technique to use more complex graph neural network architectures, such as graph transformers. Future work should also address the wide variety of preprocessing steps that could be performed on LAMP features. Due to their nature, LAMP dimensions roughly follow a power-law distribution with many dimensions close to or equal to zero and with very few extremely large values. One simple way to preprocess these features would be to take the natural log of each dimension prior to use, which would have the effect of pulling these feature distributions closer to a linear distribution, which would be more suitable for neural network input. Recent literature also suggests that evaluating the models with balanced and unbalanced datasets may reveal further avenues of improvement^27^.

Edge imputation remains a critical problem in the field of gene interaction networks. We believe that motifs and subgraph search will be a critical tool in the future of this field, and that our work is a strong step in that direction. We invite the community to use our code and data to further explore this exciting area of research, and we welcome feedback and collaboration with researchers in the field.

## Acknowledgements

We gratefully acknowledge the support of Johns Hopkins University Applied Physics Laboratory internal research and development funding which enabled this work.

## Code and data availability

All code produced for this study will be made available upon publication. DotMotif is available at https://github.com/aplbrain/dotmotif. A reference implementation of GrandIso is available at https://github.com/aplbrain/grandiso-networkx.

## Notes

### Competing Interest Statement

The authors have declared no competing interest.

